## Supplementary Figures for "Prions induce minor genome-wide translational changes in neurons compared to glia"

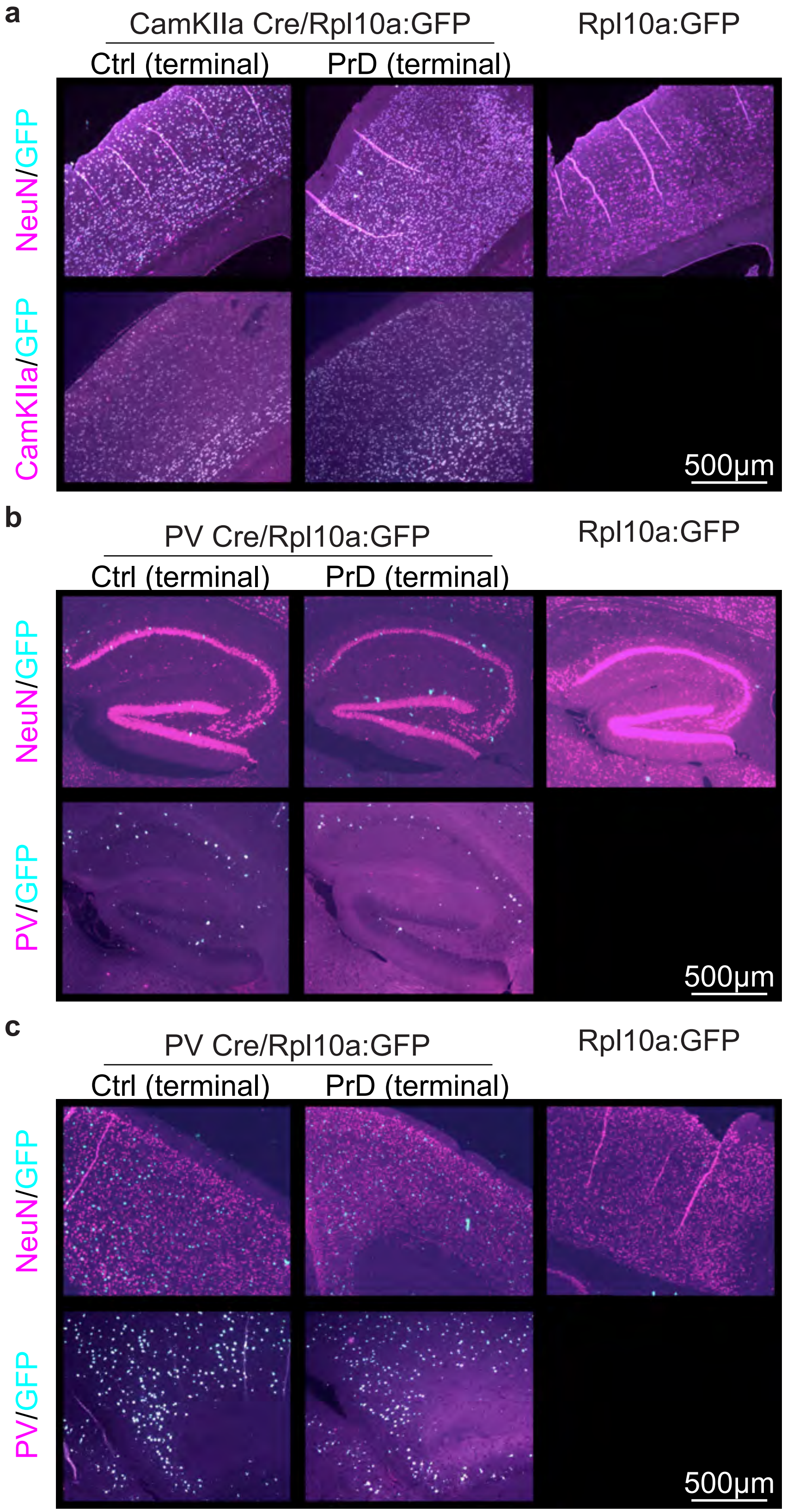


Scheckel et al., Supplementary Figure 1

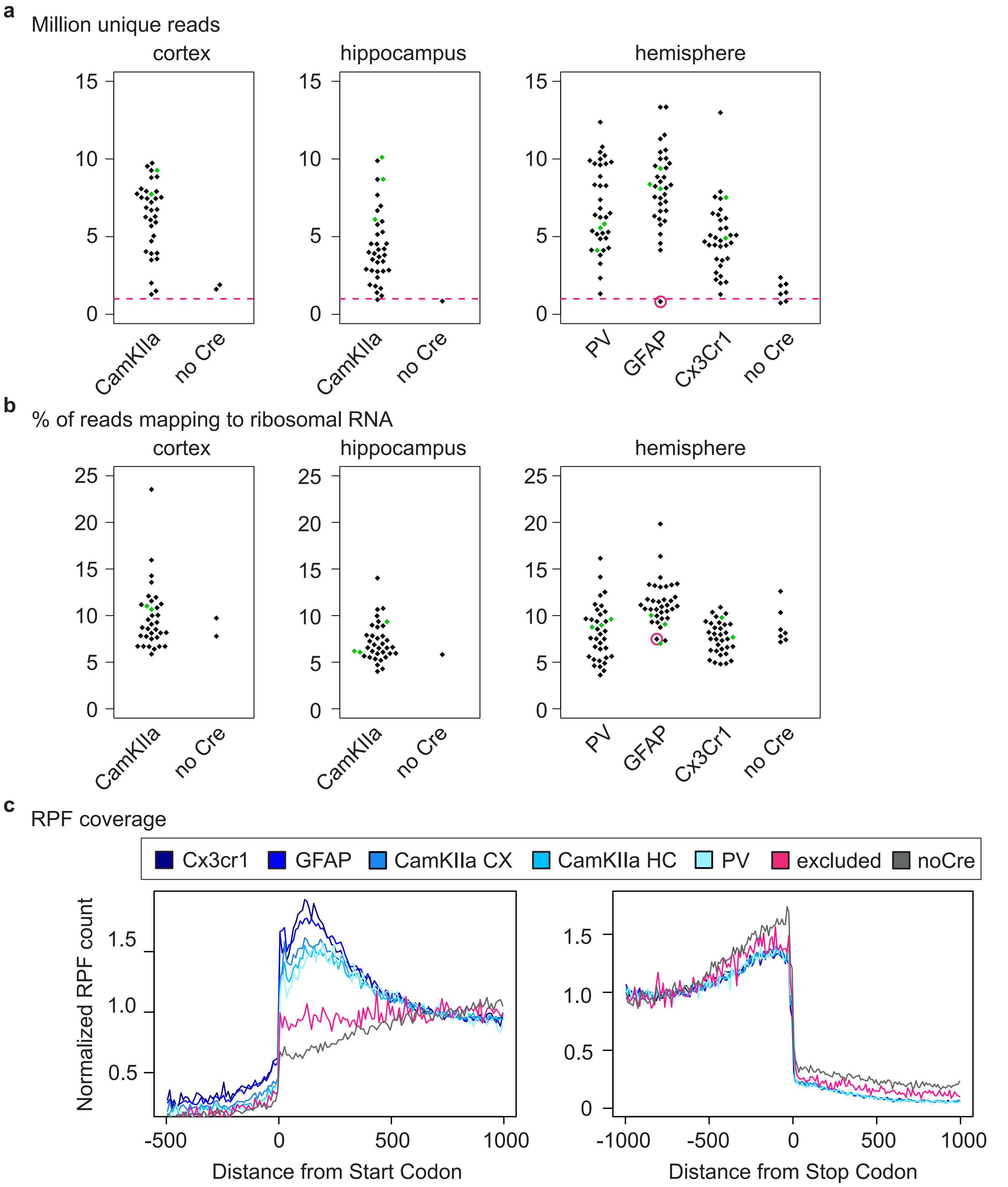


Scheckel et al., Supplementary Figure 2

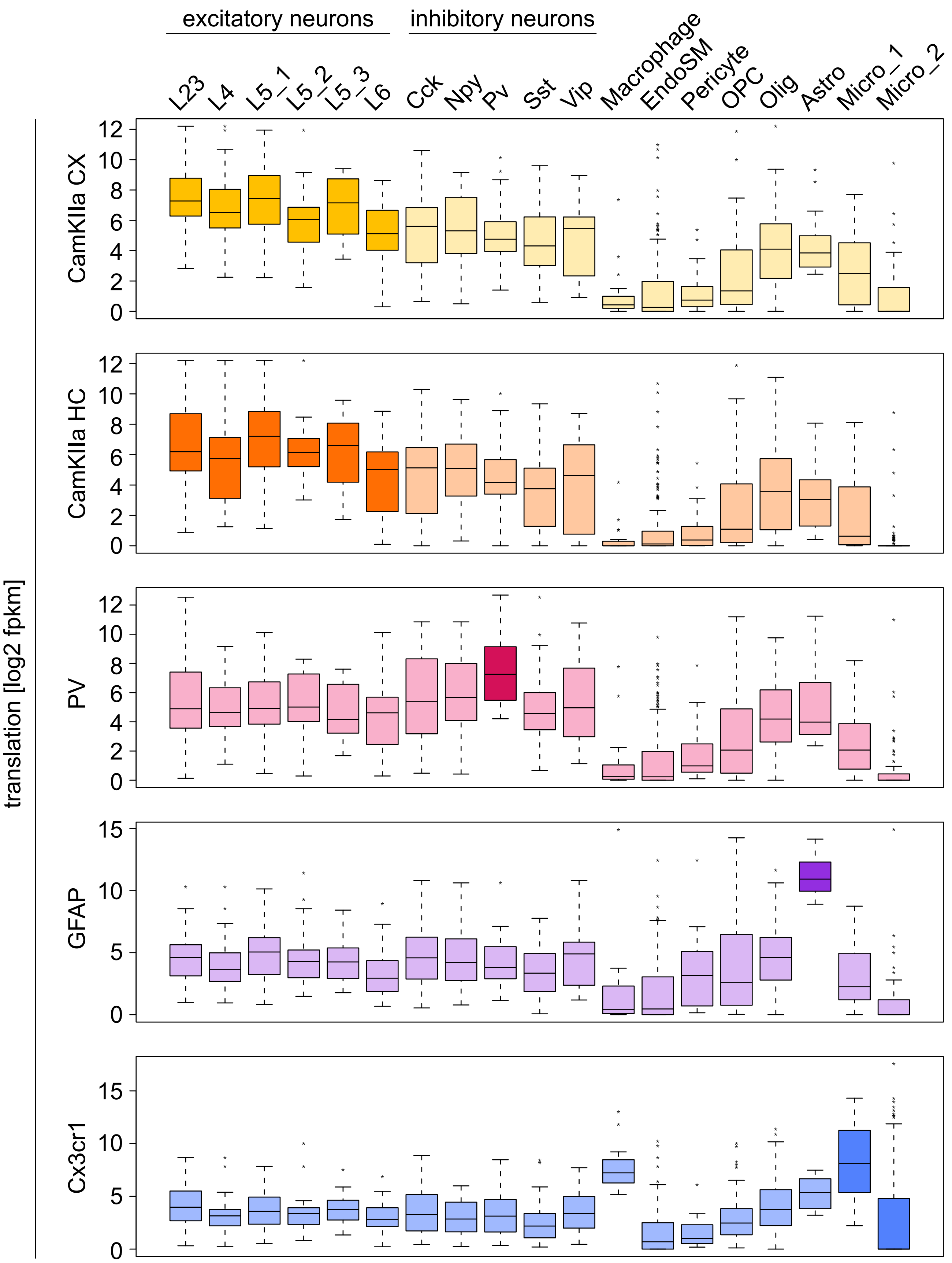


Scheckel et al., Supplementary Figure 3

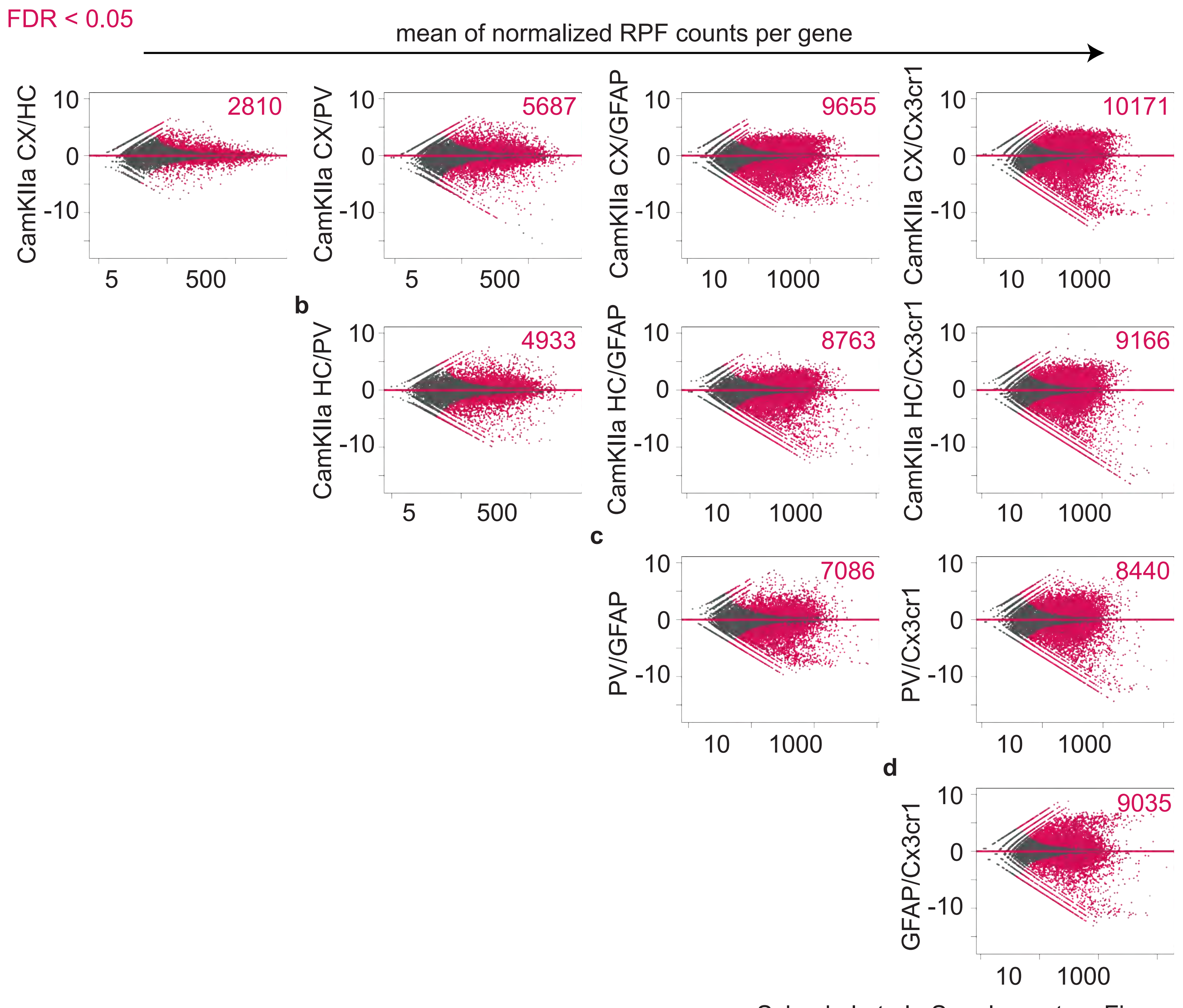


Scheckel et al., Supplementary Figure 4

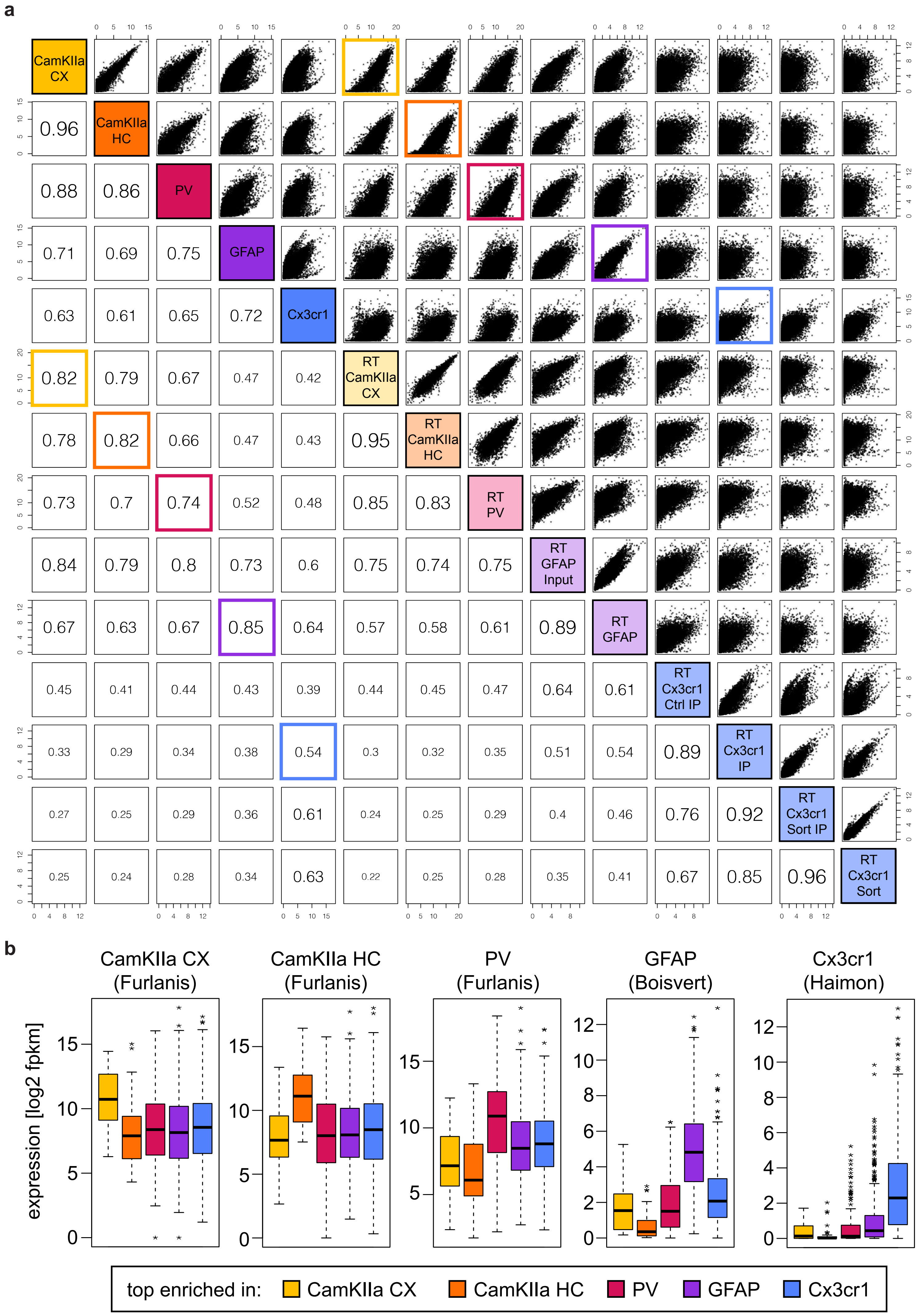


Scheckel et al., Supplementary Figure 5

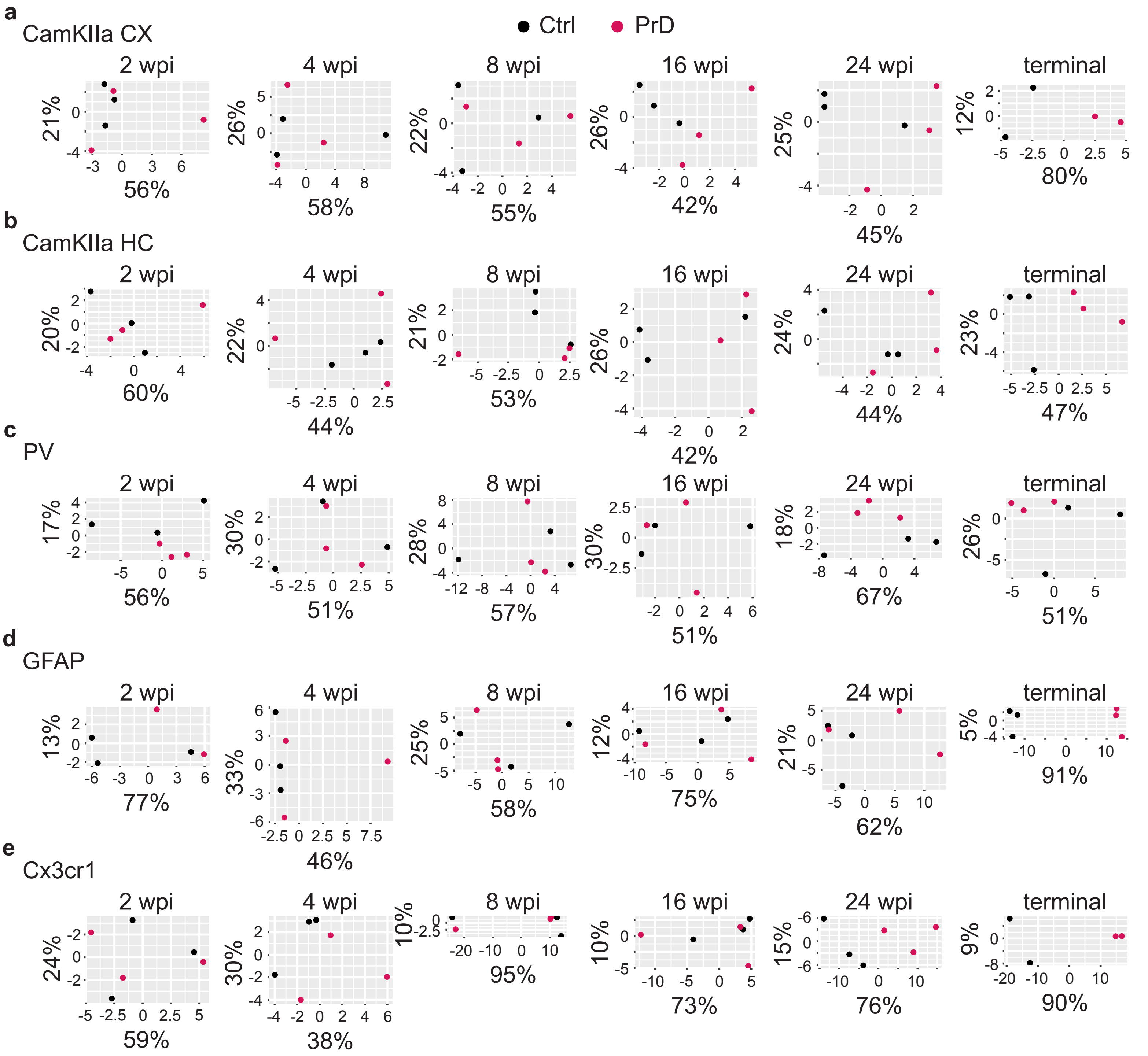


Scheckel et al., Supplementary Figure 6

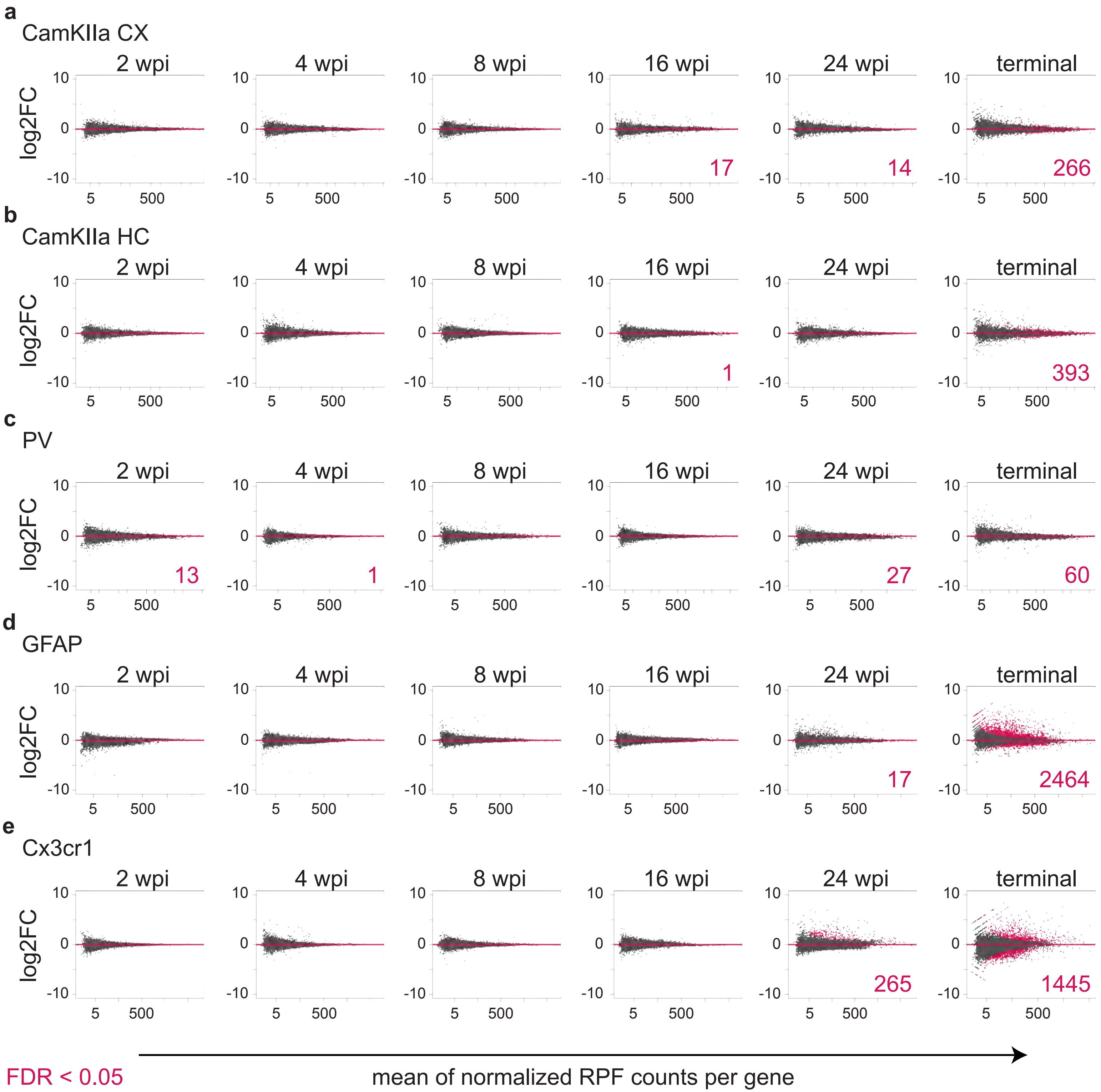
a
CamKIIa CX
2 wpi
4 wpi
8 wpi
16 wpi
24 wpi
terminal
log2FC
10
0
-10
5
500
17
14
266
b
CamKIIa HC
2 wpi
4 wpi
8 wpi
16 wpi
24 wpi
terminal
1
393
c
PV
2 wpi
4 wpi
8 wpi
16 wpi
24 wpi
terminal
13
1
27
60
d
GFAP
2 wpi
4 wpi
8 wpi
16 wpi
24 wpi
terminal
17
2464
e
Cx3cr1
2 wpi
4 wpi
8 wpi
16 wpi
24 wpi
terminal
265
1445
FDR < 0.05
mean of normalized RPF counts per gene

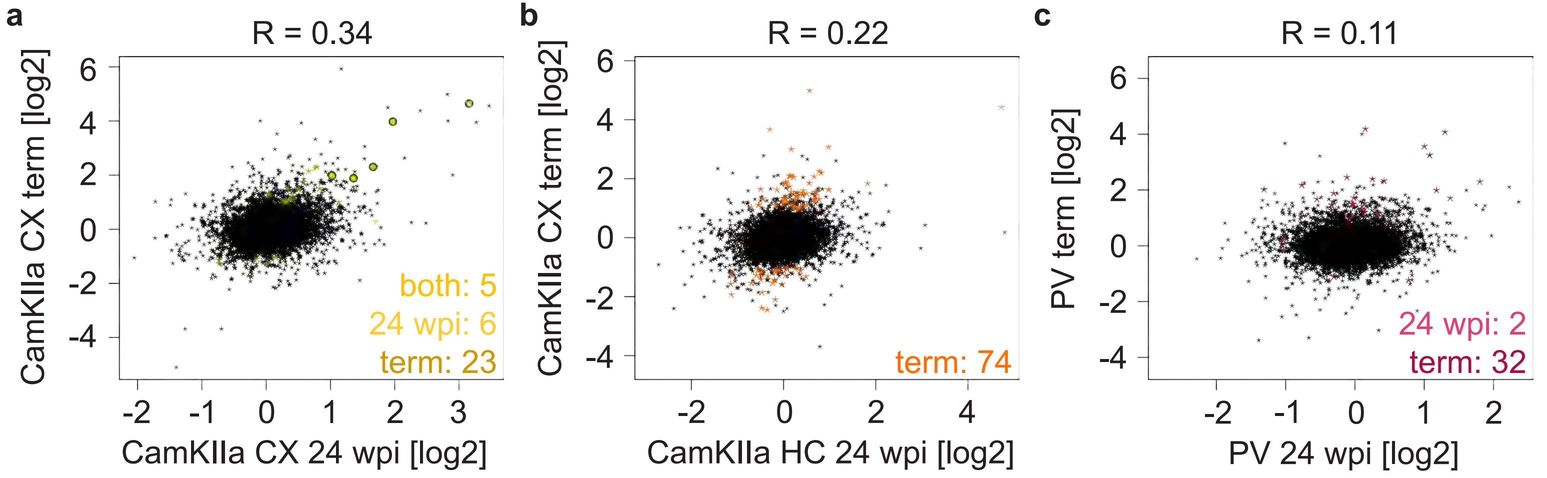


Scheckel et al., Supplementary Figure 8

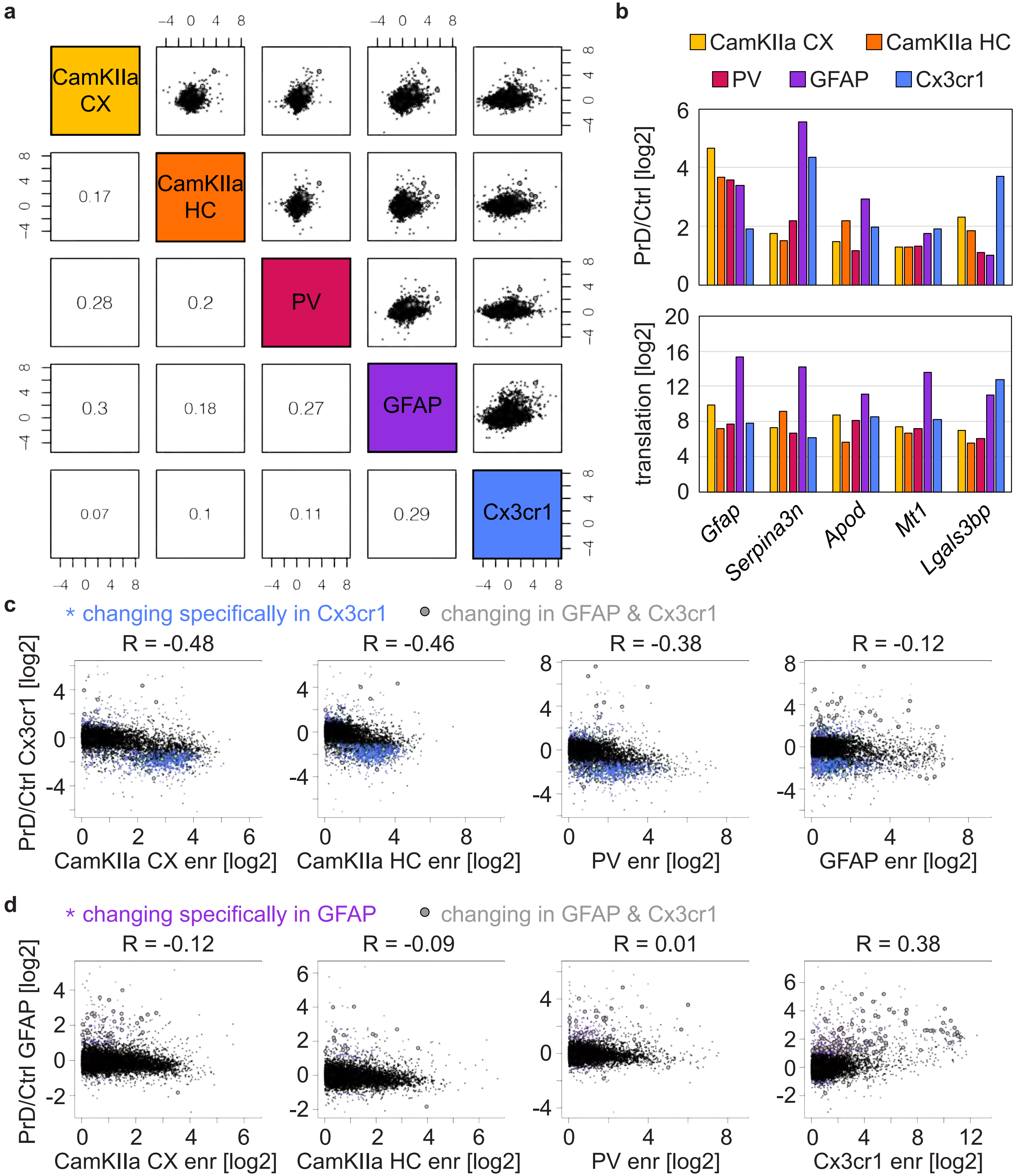


Scheckel et al., Supplementary Figure 9

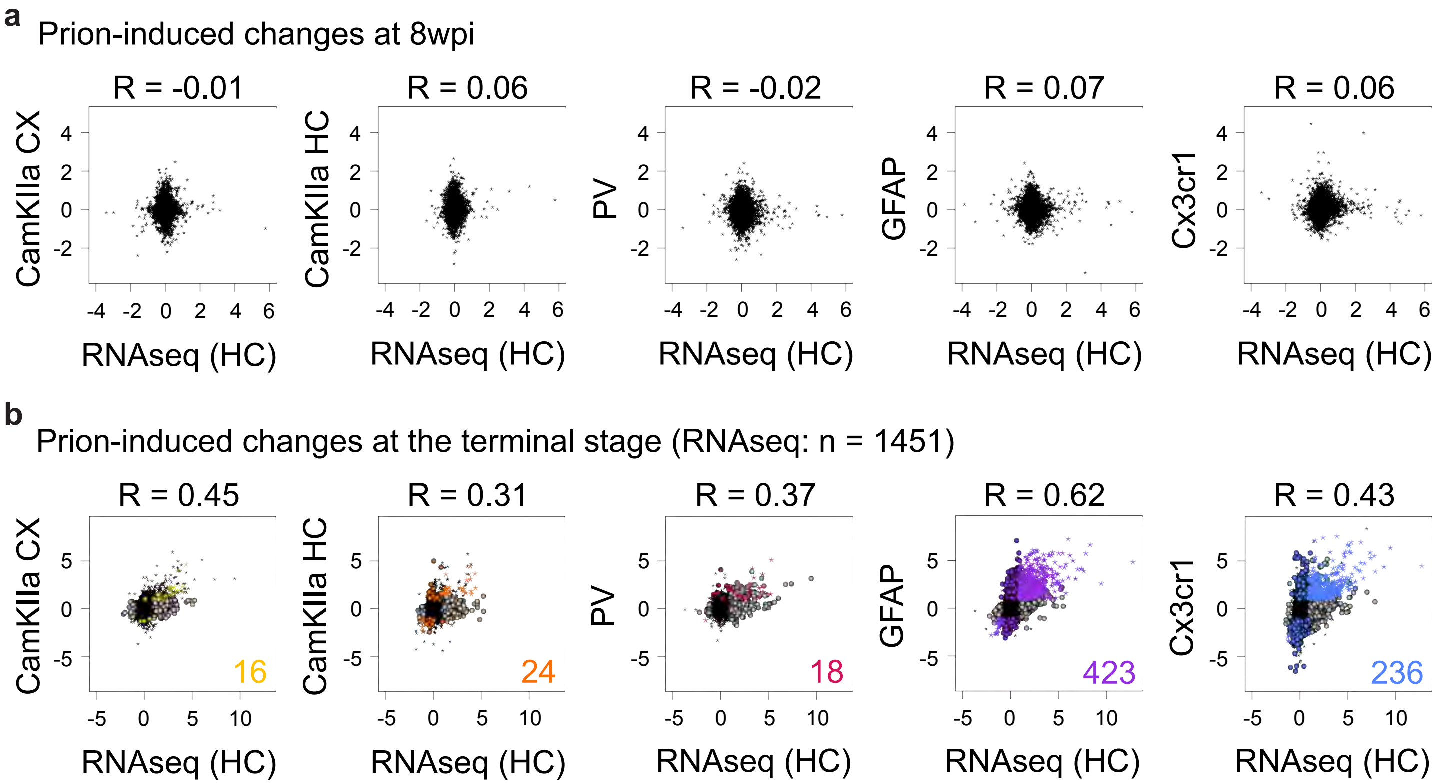


Scheckel et al., Supplementary Figure 10
